# A Covalent Organic Framework-Inspired β-Ketoenamine Crosslinking Strategy for Robust, Injectable Bovine Serum Albumin Hydrogels with pH-Triggered Drug Release

**DOI:** 10.64898/2026.09.06.749658

**Authors:** Sonal Khaitan, Tanya Agrawal, Parth Gulati, Akansha, Natasha, Tatini Rakshit, Suchetan Pal

## Abstract

Globular proteins are difficult to convert into robust hydrogels, as their compact, folded structures bury reactive residues, forcing conventional strategies to rely on denaturation or synthetic-polymer reinforcement that compromise the native protein. Inspired by the β-ketoenamine bond-forming chemistry of covalent organic frameworks (COFs), we report the crosslinking of native bovine serum albumin (BSA) with 1,3,5-triformylphloroglucinol (TFP), a C3-symmetric trialdehyde, into a chemically defined hydrogel. TFP reacts with surface-exposed lysine residues through an irreversible enol-to-keto tautomerization, confirmed by FTIR and NMR spectroscopy, generating stable β-ketoenamine crosslinks under mild aqueous conditions without denaturing the protein, as verified by intrinsic tryptophan fluorescence. The resulting hydrogels are mechanically robust compared to a reversible-imine control, injectable and self-recovering, exhibit reversible shape memory and substantial load-bearing capacity, and remain stable across a broad pH range over extended periods. The network shows consistent swelling behavior at physiological and mildly acidic pH, with modest compaction under strongly basic conditions; scanning electron microscopy reveals a dense, nodular network for the TFP hydrogel versus an open, sheet-like lamellar morphology for the reversible-imine control. The hydrogel efficiently encapsulates doxorubicin and displays pH-triggered, acid-selective release, which comparative kinetic analysis attributes principally to pH-dependent weakening of DOX–BSA binding affinity (linked to the N-to-F conformational transition of BSA near its isoelectric point) rather than to bulk network swelling or degradation. Doxorubicin-loaded hydrogels show enhanced killing of MCF-7 breast cancer cells relative to the free drug while remaining cytocompatible toward normal mammalian cells, and a ciprofloxacin-loaded variant exhibits potent antibacterial activity against both Gram-positive (M. luteus) and Gram-negative (E. coli) bacteria. This work translates reticular β-ketoenamine chemistry into a general platform for robust, stimuli-responsive protein biomaterials.

**TABLE OF CONTENTS GRAPHIC:** 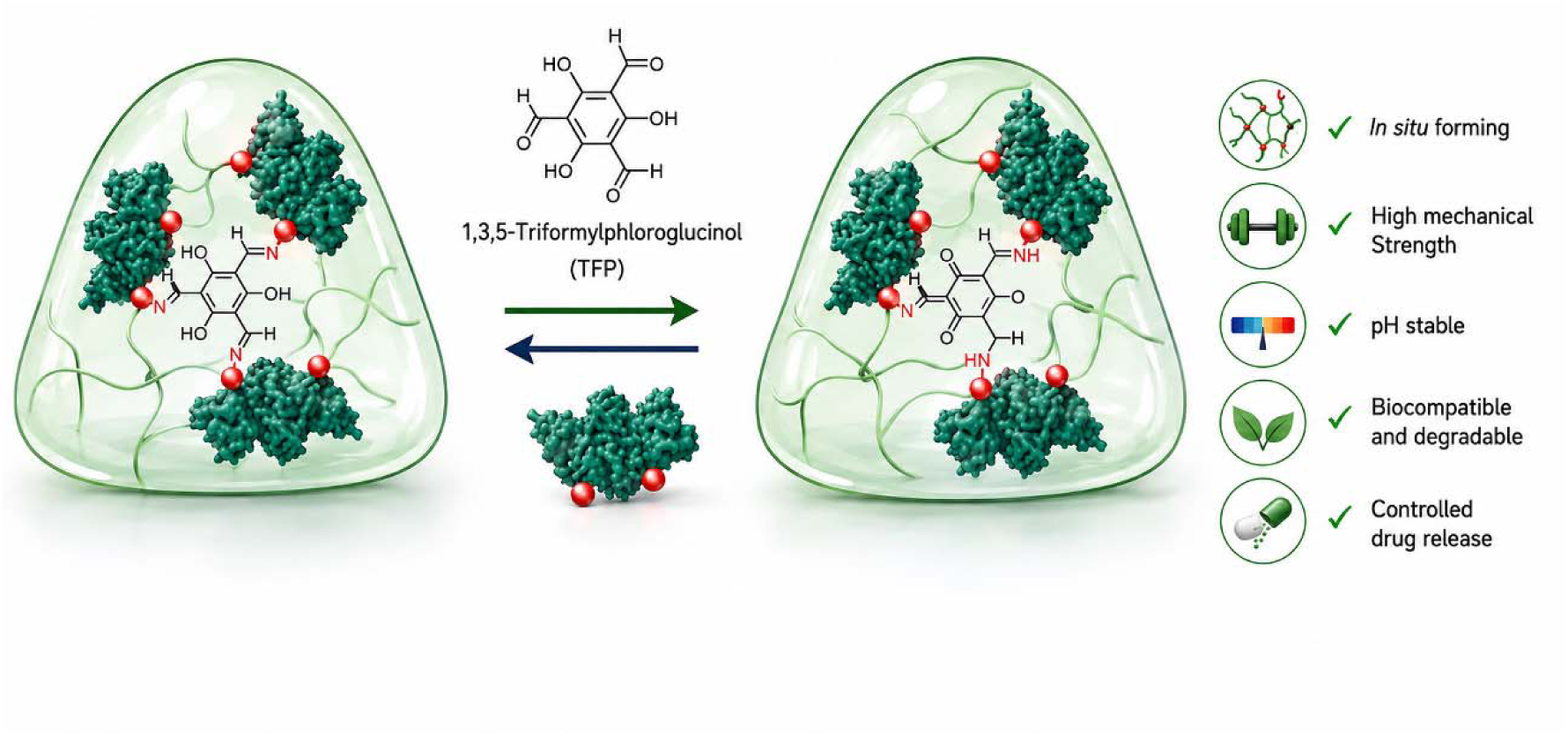

## INTRODUCTION

Protein-based hydrogels have emerged as a compelling class of biomaterials owing to their intrinsic biocompatibility, biodegradability, and rich surface chemistry.^1–3^ To date, the majority have been constructed from fibrillar proteins such as collagen,^4^ silk fibroin,^5^ fibrin,^6^ and keratin,^7^ whose extended backbone structures naturally facilitate self-assembly and network formation into mechanically robust three-dimensional matrices.^8,9^ In contrast, globular proteins pose a significant fabrication challenge: their compact, tightly folded tertiary structures bury reactive residues within a hydrophobic core, rendering conventional crosslinking strategies ineffective under mild conditions.^13–15^ Existing approaches have relied on heat-induced denaturation to expose buried reactive groups^13^ or on double-network architectures incorporating synthetic polymers,^16,17^ strategies that inevitably compromise protein structural integrity, introduce cytotoxic components, or add synthetic complexity that undermines biological utility.^13,16,18^

Bovine serum albumin (BSA), the most abundant plasma protein, has been widely explored as a hydrogel building block for drug delivery, wound healing, and tissue engineering owing to its low cost, biocompatibility, non-immunogenicity, and abundant functional residues.^10,11^ A range of gelation strategies has been reported, including physical self-assembly, thiol–disulfide exchange, photo-crosslinking, and chemical crosslinking with mono- and di-aldehydes such as glutaraldehyde and glyoxal.^11,12^ However, aldehyde-based crosslinking of BSA generally proceeds through reversible Schiff-base (imine) linkages that are hydrolytically labile, limiting mechanical and pH stability, whereas harsher covalent routes frequently require partial denaturation of the protein.^12^ A crosslinking chemistry that forms hydrolytically stable, irreversible bonds directly on the native, folded protein under mild aqueous conditions and without added polymers would therefore represent a meaningful advance for albumin-based biomaterials.

More broadly, the chemical crosslinking of proteins is dominated by an established but limited toolkit: glutaraldehyde,^19,20^ carbodiimide (EDC/NHS) coupling,^21^ genipin,^22,23^ transglutaminase,^24^ and SpyCatcher–SpyTag chemistry^25,26^ each carrying well-documented limitations of toxicity, reversibility, or biological complexity (Figure 1). We reasoned that a fundamentally different bond-forming chemistry, drawn from reticular materials science, could address these limitations.

**Figure 1.**
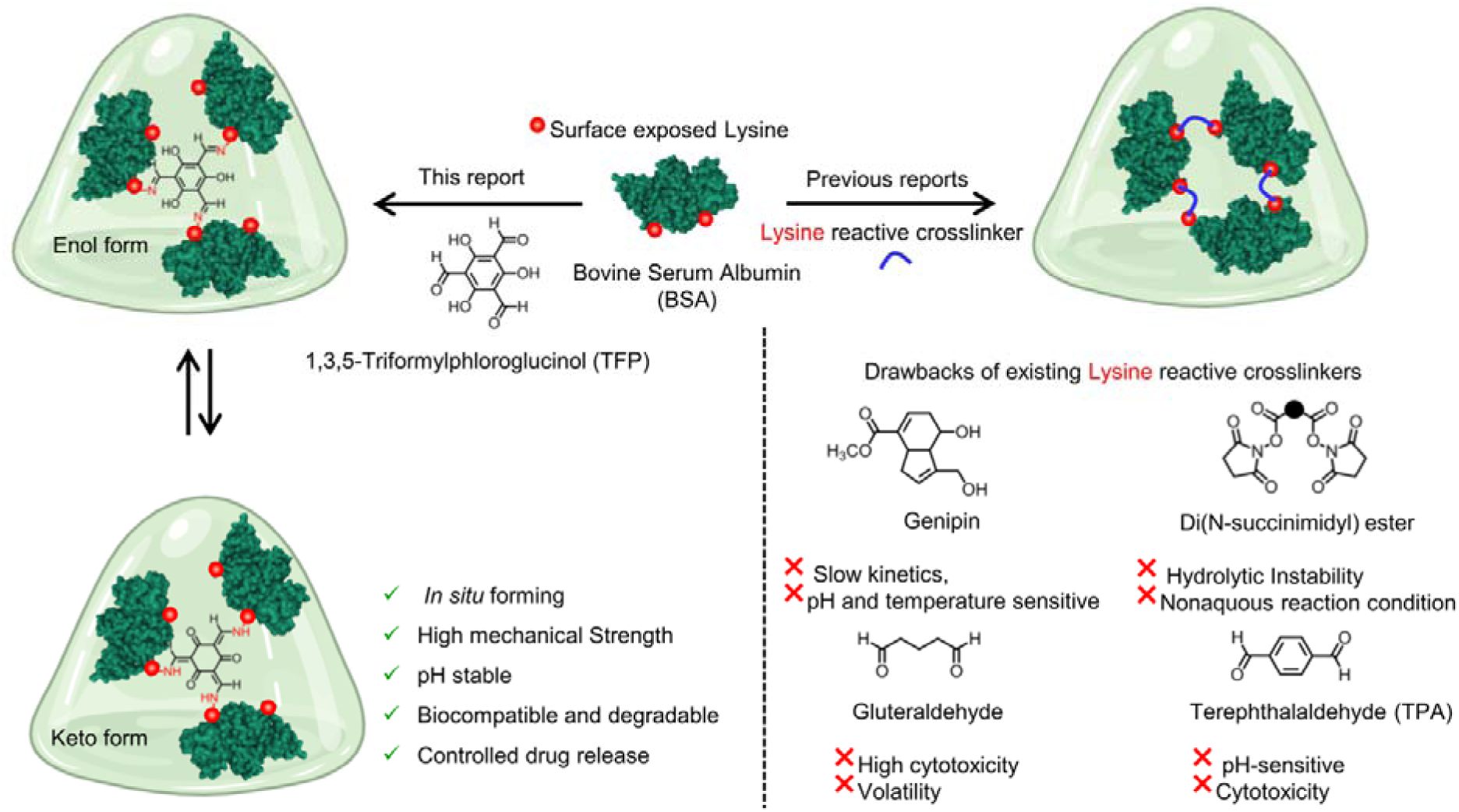
Schematic illustration of β-ketoenamine crosslinking of BSA for fabricating a chemically crosslinked globular protein hydrogel. Left: TFP exists in a keto–enol tautomeric equilibrium and reacts with surface-exposed lysine residues of BSA to form β-ketoenamine crosslinks, yielding a mechanically robust, *in situ* forming, pH-stable, biocompatible, and biodegradable hydrogel with controlled drug-release capability. Right: comparison with previously reported lysine-reactive crosslinkers.

In the field of covalent organic frameworks (COFs), 1,3,5-triformylphloroglucinol (TFP), a C_3_-symmetric trialdehyde, reacts with primary amines through a combination of reversible Schiff-base condensation and an irreversible enol-to-keto tautomerization to generate exceptionally stable β-ketoenamine linkages.^27,28^ This chemistry endows keto-enamine COFs with outstanding chemical and hydrolytic stability. We hypothesized that the same irreversible β-ketoenamine bond formation could be harnessed to crosslink the surface lysine residues of a native globular protein, translating a design principle from crystalline, reticular solids to soft, functional biomaterials.

Herein, we report the first application of TFP as a trifunctional crosslinker for a native globular protein, using BSA as the model. TFP crosslinks surface-exposed lysine residues under mild aqueous conditions, enabling rapid, catalyst-free gelation into a robust, chemically stable network without protein denaturation.^29–32^ We establish the β-ketoenamine crosslinking chemistry spectroscopically, characterize the network morphology, swelling behavior, and mechanical properties (including under equilibrium-swollen conditions), and demonstrate pH-triggered doxorubicin delivery with enhanced anticancer efficacy and cytocompatibility toward normal cells. Together, these results position irreversible β-ketoenamine chemistry as a general route to mechanically robust, stimuli-responsive protein biomaterials.

## EXPERIMENTAL SECTION

### Materials

Bovine serum albumin (BSA), doxorubicin hydrochloride (DOX), and L-lysine were obtained from TCI Chemicals. 1,3,5-Triformylphloroglucinol (TFP) and terephthaldehyde (TPA) were obtained from BLD Pharma and used without further purification. Proteinase K (from Tritirachium album) was obtained from Sigma-Aldrich. MCF-7 human breast cancer cells were obtained from the National Centre for Cell Science (NCCS), Pune, India. High-glucose Dulbecco’s modified Eagle’s medium (DMEM), fetal bovine serum (FBS), and penicillin–streptomycin were obtained from Gibco. All other reagents and solvents were of analytical grade and used as received. The gelation buffer was 500 mM potassium carbonate buffer (pH ∼9.5).

### Preparation of TFP- and TPA-Crosslinked BSA Hydrogels

BSA and the crosslinker (TFP or TPA) were separately dissolved in 500 mM carbonate buffer (pH ∼9.5) supplemented with L-lysine to improve crosslinker solubility. The two solutions were mixed in a 1:1 volume ratio to give final BSA concentrations of 200–250 mg/mL and crosslinker concentrations of 75–100 mM, and were incubated at ambient temperature. Gelation was assessed by the tube-inversion method. We removed the additional crosslinker by washing the hydrogel with PBS twice for further studies.

### Fourier-Transform Infrared (FTIR) Spectroscopy

FTIR spectra of free BSA, free lysine, TFP, TPA, and their reaction products were recorded in attenuated total reflectance (ATR) mode on a Spectrum Two FTIR spectrometer (PerkinElmer) over 400–4000 cm⁻¹. Free lysine was used as a low-molecular-weight model for the surface lysine residues of BSA.

### Nuclear Magnetic Resonance (NMR) Spectroscopy

Solution-state ¹H and ¹³C NMR spectra of free TFP and the TFP–lysine reaction product were acquired on a 600 MHz Bruker spectrometer in a mixed solvent of DMSO-d₆ (70%) and D₂O (30%).

### Intrinsic Tryptophan Fluorescence

Emission spectra were recorded on a microplate reader (Molecular Devices, SpectraMax i3x) with excitation at 295 nm. Spectra were collected over an extended emission window to ∼590 nm to probe both tryptophan emission and any β-ketoenamine-associated emission. Native BSA and thermally/chemically denatured BSA were measured as references. This 295 nm/∼590 nm scan is a separate, additional measurement to the 280 nm-excitation, 300–450 nm native-fold comparison reported and was performed specifically to probe for a distinct β-ketoenamine emission signature within the intact TFP hydrogel. Because the hydrogel is formed and equilibrated in the alkaline (pH ∼9.5) carbonate gelation buffer, the β-ketoenamine chromophore’s emission was substantially quenched under these basic conditions, and only a very low-intensity feature was observed.

### Rheological Characterization

Oscillatory rheology was performed on an Anton Paar Rheocompass rheometer with parallel-plate geometry (25 mm diameter, 1 mm gap) at room temperature. Frequency sweeps (0.1–100 rad/s) were performed within the linear viscoelastic region at a constant strain of 0.5%. Strain (amplitude) sweeps were performed at a constant angular frequency of 1 rad/s over a strain range of 0.1–1000%. Cyclic step-strain measurements alternated between high strain (500%) and low strain (0.5%) at a constant angular frequency of 1 rad/s over four consecutive cycles to assess injectability, yielding behavior, and self-recovery. Measurements were performed on both as-prepared gels and on gels equilibrated to their equilibrium-swollen state, and the hydration state is specified for each measurement.

### Equilibrium and pH-Dependent Swelling

Pre-weighed, dried hydrogel discs (dry mass, W) were immersed in the swelling medium at 37 °C and periodically blotted to remove surface liquid and weighed (W) until the mass stabilized; the equilibrium swelling ratio was calculated as Q = (W − W)/W. For the physiological-pH measurement (W = 413 mg), mass uptake plateaued, giving Q ≈ 1.17 at pH 7.4. pH-dependent swelling was measured by equilibrating gels in PBS at pH 5.5, 7.4, and 13 at 37 °C, with mass recorded at 1, 4, 8, 12, and 24 h; equilibrium was reached by 24 h at all three pH values.

### Scanning Electron Microscopy (SEM) of hydrogels

Hydrogels were first gold-coated, then mounted and imaged by secondary-electron microscopy at an accelerating voltage of 2.00 kV and a working distance of ∼15.7 mm to visualize the network morphology and pore structure of the TFP and TPA hydrogels.

### Load-Bearing and Shape-Memory Testing

Wire-shaped hydrogel specimens were fabricated by molding and subjected to increasing static tensile loads. Dehydration–rehydration cycles were performed by ambient drying and re-immersion in water, and specimen lengths were recorded to quantify reversible shape recovery.

### pH Stability

Hydrogels were incubated in PBS spanning pH 1–13 at 37 °C, and structural integrity was assessed by tube inversion and visual inspection over 168 h. Mechanical stability was evaluated by frequency-sweep rheology on gels incubated at pH 4, 5.5, 7.5, and 13 for 4 h at 37 °C.

### Enzymatic Biodegradation

Hydrogels were incubated in 1% (w/v) Proteinase K solution at 37 °C, with protease-free samples as controls, and integrity was monitored by tube inversion over seven days.

### Bacterial Inhibition Zone Test

Bacterial solution (50 µL) of 0.1 OD (10^8^ CFU/ml) was spread over the LB agar plates using an L-spreader. The hydrogel was then kept in the centre of the plate. After a day of culturing, the inhibition zone diameter was captured.

#### Scanning Electron Microscopy of Bacteria

A mortar and pestle were used to lyophilize and pulverize the HGs. A tiny quantity of it was adhered to a pristine, conductive surface. Bacterial samples were dehydrated using a graded ethanol series (30%, 50%, 70%, and 90%) for 15 minutes at each stage after being fixed with 2.5% glutaraldehyde for 0.5 hours. A Gemini 500 field emission scanning electron microscope from Zeiss, Germany, was then used to image the samples.

### In Vitro Surface Antibacterial Test

Hydrogel formulations TFP-BSA were prepared by adding various hydrogel precursors, including antibacterial precursors, to the microcentrifuge tube. Once the gels were formed, a bacterial suspension of 10^8^ CFU/ml (0.1 OD) was carefully poured over the surface of the gels. The microcentrifuge tube was incubated for 3 hours at 37°C in a BOD incubator. Subsequently, 500µl of sterile 1x PBS was added over the treated bacteria to resuspend any surviving bacteria. Control was prepared by directly mixing 500 μL of sterile 1x PBS with bacterial suspension of 10^8^ CFU/ml concentration (0.1 OD). 300µl of treated bacteria suspension was collected from the surface of the gels and washed twice with sterile 1x PBS. Finally, 50µl bacterial suspension was spread on the agar plate. After 12-16 hrs, bacterial growth was assessed to evaluate the antibacterial efficiency of the loaded TFP-BSA.

### Bacterial Cell Culture source and Preparation

*E.coli (MTCC No. 11948) and M.luteus (MTCC No. 11948) were procured from the Microbial Type Culture Collection* and Gene Bank (MTCC), Chandigarh. Bacterial cultures of *E.coli* (Gram-negative) and *M.luteus* (Gram-positive) were prepared using Luria-Bertani (LB) broth. 2g of LB powder was dissolved in 100 ml of distilled water and subsequently autoclaved at 121°C at 15 psi to ensure proper sterilization. Once cooled, the broth is transferred aseptically to separate Falcon tubes. Subsequently, 50 µl of the bacteria is transferred to each tube and incubated overnight at 37°C at 180 rpm on a rotary shaker incubator. Bacterial growth is confirmed by observing turbidity in the tubes. For subculturing, the overnight culture was diluted in 1:100 dilution for 3 hrs, following the same conditions as of overnight culture until the OD 600 reached approximately 1.0, indicating the logarithmic growth phase.

### Drug Loading and Loading Capacity

Ciprofloxacin was loaded during gelation.100 ul of 0.1mg/ml ciprofloxacin was taken and mixed with the precursor solutions. DOX was loaded in situ by dissolving it in the precursor solution before gelation. After gelation and removal of non-encapsulated drug, the encapsulated DOX was quantified by intrinsic fluorescence against a calibration curve constructed from free-DOX standards (0.5, 1, 1.5, 1.75, and 2 mg/mL), giving the linear regression y = 2,924,095.5x + 4,737,304.4. Hydrogels prepared with nominal DOX loading concentrations of 0.5, 1.0, 1.5, and 1.75 mg/mL were measured by the same method, and the true encapsulated DOX concentration was back-calculated from this equation. Drug loading efficiency (DLE, %) was calculated as (actual encapsulated DOX concentration / nominal DOX concentration added) × 100.

### In Vitro Drug Release and Release Kinetics

Drug release of two drug molecules was carried out, namely Ciprofloxacin and Doxorubicin. Cipro-loaded hydrogels were in a buffer at pH 7.4, while DOX-loaded hydrogels were immersed in buffer at pH 4, 7.4, and 10 at 37 °C. At each timepoint (1, 2, 3, 4, 6, 12, 24, 36, and 48 h), a 100 μL aliquot of the release medium was withdrawn in triplicate from 1 mL of loaded hydrogel incubated with 10 mL of 10 mM PBS in an orbital shaking incubator, and released DOX was quantified by intrinsic fluorescence (excitation 470 nm), ciprofloxacin was quantified in parallel aliquots 270 nm on a microplate reader (Molecular Devices, SpectraMax i3x) against a standard calibration curve. Rather than fitting the release profiles to a diffusion-based model (e.g., Korsmeyer–Peppas), the release mechanism was established by comparative kinetics analysis: (i) the timescale of enzymatic network degradation (days, and protease-dependent; see Enzymatic Biodegradation) and the timescale of structural integrity loss at extreme pH (>24 h, and only outside pH 4–10) were both far slower than the 24–48 h window over which DOX release occurred, excluding bulk degradation as the driver; (ii) the equilibrium swelling ratio changed only marginally between pH 5.5 (Q = 1.19) and pH 7.4 (Q = 1.17) far too small a difference to account for the 2.8-fold difference in cumulative release between pH 4 and pH 7.4 at 24 h excluding bulk swelling as the dominant driver. Release is therefore attributed principally to a pH-dependent decrease in Drug–BSA binding affinity, with a minor, secondary contribution from acid-enhanced network swelling.

### Cell Culture and Cytocompatibility

MCF-7 human breast cancer cells and HEK 293 cells were cultured in high-glucose DMEM supplemented with 10% FBS and 1% penicillin–streptomycin at 37 °C in a 5% CO₂ atmosphere; cells at passages 3–5 were used for all experiments. Cells were seeded in 96-well plates at a density of 1×10 cells per well and allowed to adhere for 24 h before treatment with the TFP hydrogel at 0.01, 0.1, 1, and 100 mg/mL for 24 h. Cell viability was assessed using the WST-1 reagent (colorimetric assay, Invitrogen) according to the manufacturer’s instructions, with absorbance recorded at 450 nm.

### Apoptosis Analysis by Flow Cytometry

MCF-7 cells were seeded in 12-well plates at 1×10 cells per well and allowed to adhere for 24 h. Cells were treated with free DOX (1.5 μg/mL) or DOX-loaded hydrogels (DOX loading 0, 5, 10, and 20 mg/mL) for 6 h, then harvested, washed with PBS, and stained with Annexin V/PI (Elabscience) according to the manufacturer’s instructions. Flow cytometry was performed on a Beckman Coulter CytoFLEX S instrument, acquiring a minimum of 10,000 events per sample; data were analyzed using CytoFLEX S software.

### Statistical Analysis

Data are expressed as mean ± standard deviation (n = 3 unless otherwise stated). Statistical significance was assessed by *t-test,* with p < 0.05 considered significant.

## RESULTS AND DISCUSSION

### Design and Fabrication of TFP-Crosslinked BSA Hydrogels

As a model globular protein, BSA possesses 30–35 surface-accessible lysine residues (out of 59 total), each bearing a primary ε-amine well suited to reaction with the trialdehyde TFP (Figure 1).^33^ We used terephthaldehyde (TPA), a bifunctional dialdehyde that forms only reversible imine linkages, as a mechanistic control. BSA and TFP (or TPA) solutions were mixed in a 1:1 volume ratio in 500 mM carbonate buffer supplemented with lysine to improve solubility. Stable hydrogels formed within 3 h under ambient conditions, as judged by tube inversion, exclusively at crosslinker concentrations of 75 and 100 mM combined with BSA at 200 and 250 mg/mL (Figure 2a, b), defining a well-bounded gelation window.

**Figure 2.**
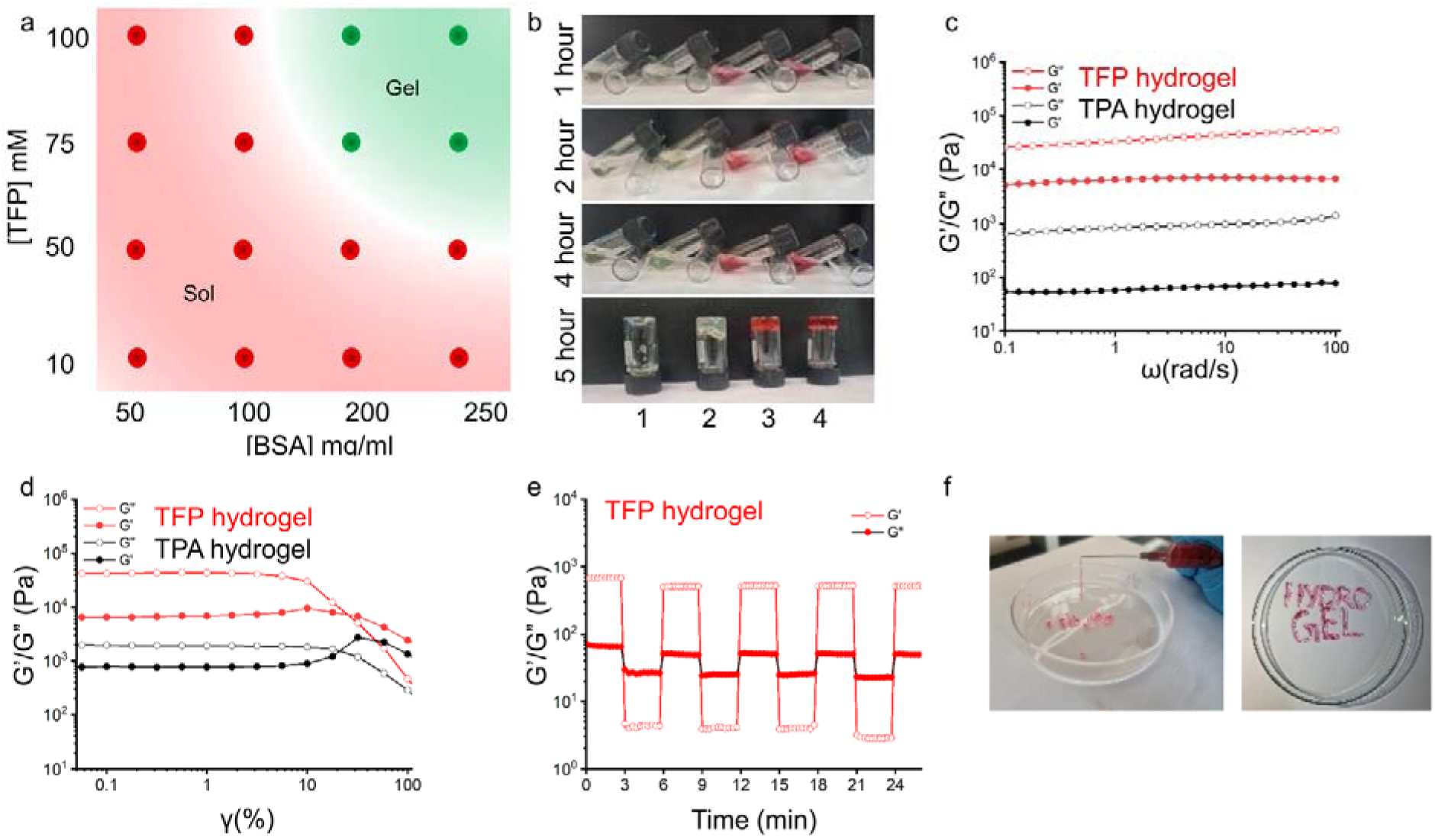
(a) Phase diagram mapping the gelation window as a function of TFP concentration (10–100 mM) and BSA concentration (50–250 mg/mL); green and red circles denote gel and sol states, respectively. (b) Time-dependent gelation (1, 2, 4, and 5 h) for four crosslinker conditions: (1) no crosslinker, (2) TPA, (3) 50% TPA + 50% TFP, and (4) TFP. (c) Frequency sweep and (d) amplitude (strain) sweep of the TFP (red) and TPA (black) hydrogels, showing G′ > G″. (e) Step-strain measurements of the TFP hydrogel showing a reversible gel-to-sol transition under high strain and rapid recovery at low strain. (f) Photographs demonstrating the processability of the TFP hydrogel.

### β-Ketoenamine Crosslinking Chemistry

We elucidated the crosslinking chemistry of TFP and TPA with lysine by comparing FTIR spectra, using free lysine as a model for the surface lysine residues of BSA (Figure S1). Direct FTIR analysis of the intact BSA hydrogel is complicated by the O–H bending and Amide I (1650 cm^−1^) and Amide II (1540 cm^−1^) bands of the protein backbone, which overlap with crosslinker-derived peaks in the 1550–1700 cm^−1^ region (Figure S2). Free TPA exhibited a Fermi-resonance doublet at 2865 and 2805 cm^−1^ (aldehyde C–H) alongside the aromatic aldehyde C=O at 1690 cm^−1^. Free TFP showed peaks at 2890 cm^−1^ (aldehyde C–H), 1635 cm^−1^ (aldehyde C=O, keto tautomer), 1586 cm^−1^ (C=C, enol tautomer), and 1247 cm^−1^ (enol C–OH), confirming its keto–enol equilibrium. Upon reaction with lysine, both aldehydes were fully consumed. Critically, the two crosslinkers produced markedly different products: TPA + lysine showed a new imine C=N peak at 1640 cm^−1^ with broad N–H/O–H absorption at 3200 cm^−1^ (reversible Schiff base), whereas TFP + lysine showed no imine C=N and instead a broad enamine N–H at 3168 cm^−1^, a broadened β-ketoenamine C=O near 1586 cm^−1^, and a strong C–N stretch at 1359 cm^−1^ the diagnostic signatures of irreversible β-ketoenamine formation. These results show that, whereas TPA forms only a reversible imine, TFP undergoes irreversible enol-to-keto tautomerization to generate stable β-ketoenamine crosslinks, directly analogous to those in TFP-based COFs and demonstrated here for the first time in a protein-crosslinking context.

Solution-state NMR corroborated the keto-enol equilibrium of free TFP (Figure S3). The ^1^H spectrum showed three peaks at δ 9.5, 8.3, and 7.7 ppm, assignable to the aldehyde C–H (keto), enol O–H, and olefinic C–H (enol form), and the ^13^C spectrum showed two peaks in the 150–200 ppm region for the aldehyde carbonyl (keto) and enol carbons. Upon reaction with lysine, the equilibrium collapsed into a single ^13^C peak at 168 ppm, distinct from the free aldehyde (∼190 ppm) and a typical imine carbon (∼158–162 ppm), consistent with β-ketoenamine formation and complementing the FTIR data.

### Preservation of the Native Protein Fold

To confirm that crosslinking does not denature BSA, we performed intrinsic tryptophan fluorescence spectroscopy (Figure S4). The TFP hydrogel exhibited an emission maximum at 355 nm, compared with 350 nm for native BSA and 390 nm for fully denatured BSA. The minimal 5 nm red shift, in contrast to the 40 nm shift on full denaturation, indicates that TFP crosslinks surface lysine residues without perturbing the native tertiary fold. In a separate scan extending the emission window to ∼590 nm under 295 nm excitation, we looked for a distinct β-ketoenamine emission signature within the intact hydrogel; only a very weak peak was detected, most likely because the basic pH of the as-formed hydrogel (gelation buffer, pH ∼9.5) quenches keto-enamine/enaminone fluorescence, whose emissive tautomer is disfavoured under alkaline conditions. We therefore do not rely on this measurement as independent evidence of crosslink formation; FTIR and NMR (Figures S1–S3) remain our primary spectroscopic evidence for β-ketoenamine bond formation.

### Network Morphology

The internal architecture of the hydrogel was examined by scanning electron microscopy (Figure S5). The TFP hydrogel displayed a dense, compact, nodular/granular microstructure with fine, tightly packed features, consistent with a highly crosslinked, mechanically robust network. In contrast, the TPA hydrogel displayed an open, sheet-like, lamellar morphology with larger interlamellar voids, consistent with its markedly lower storage modulus and the more labile, reversible nature of its bifunctional imine crosslinks.

### Swelling Behaviour and pH Dependence

The equilibrium (gravimetric) swelling ratio of the TFP hydrogel was determined as Q = (W⍰− W□)/W□ for dried hydrogel discs (W□= 413 mg) immersed in PBS at pH 7.4, 37 °C. Mass uptake plateaued at Q ≈ 1.17 (i.e., W⍰/W□ ≈ 2.17), corresponding to water uptake equal to roughly 117% of the dry mass at equilibrium under physiological conditions (Table S1). Swelling was further measured across pH 5.5, 7.4, and 13, with equilibrium reached by 24 h at each pH (Table S1). The equilibrium swelling ratios were comparable at pH 5.5 (Q = 1.19, ∼119%) and pH 7.4 (Q = 1.17, ∼117%) and modestly reduced at pH 13 (Q = 0.97, ∼97%). This trend is consistent with the rheological data (Figure 3d), where the elevated storage modulus at pH 13 was attributed to increased crosslink density from partial BSA unfolding and exposure of additional lysine residues; a denser covalent network is expected to accommodate less water, explaining the lower equilibrium swelling ratio at this pH despite the higher stiffness. As discussed below, the near-identical swelling at pH 5.5 and 7.4 rules out bulk swelling as the principal driver of the pH-selective DOX release observed between these conditions.

**Figure 3.**
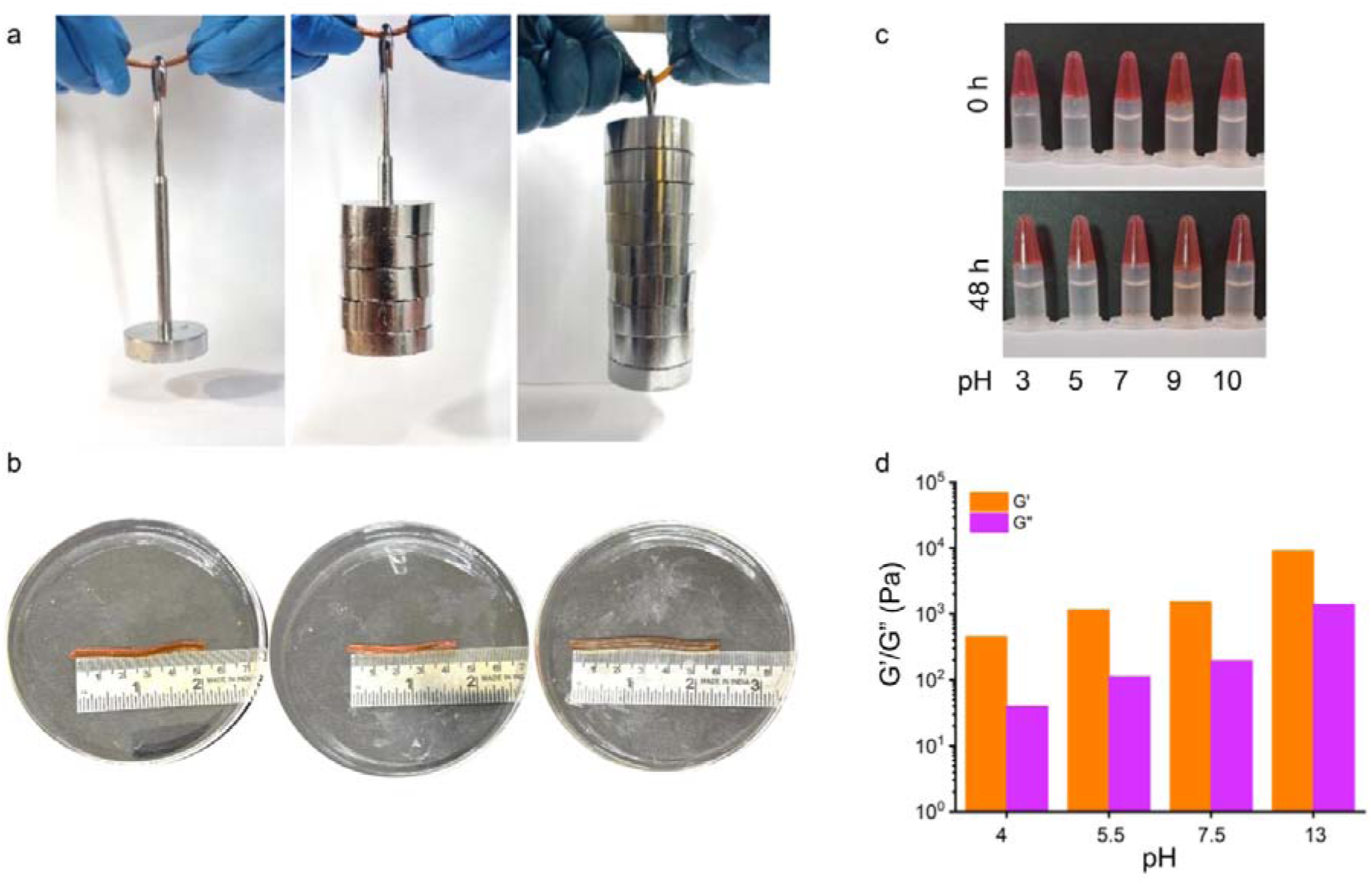
(a) Photographs demonstrating the load-bearing capacity of a TFP hydrogel wire, sustaining up to 900 g without fracture. (b) Reversible shape recovery of the TFP hydrogel wire: hydrated (∼5.6 cm), dehydrated (∼4.4 cm; ∼21% length reduction), and rehydrated (∼5.9 cm) states. (c) Photographs of TFP hydrogels in PBS at pH 3, 5, 7, 9, and 10 at 0 and 48 h, confirming retention of gel integrity across the physiologically relevant pH range. (d) Storage (G′) and loss (G″) moduli of TFP hydrogels after 4 h incubation at pH 4, 5.5, 7.5, and 13, confirming G′ > G″ at all tested pH values.

### Mechanical Properties and Injectability

We characterized the mechanical properties and injectability of the TFP and TPA hydrogels by oscillatory rheology. In frequency sweeps (0.1–100 rad/s), both hydrogels showed G′ > G″ across the entire range, confirming viscoelastic solid-like character (Figure 2c). The TFP hydrogel exhibited G′ ≈ 33,500 Pa at 1 rad/s, approximately 40-fold higher than the TPA hydrogel (839 Pa), reflecting the superior mechanical integrity of the irreversible, trifunctional β-ketoenamine crosslinks relative to the reversible, bifunctional imine linkages of TPA. Strain sweeps revealed a TFP hydrogel linear-viscoelastic plateau of G′ = 42,000–44,000 Pa up to ∼5–10% strain, with a gel-to-sol crossover at 17–31% strain (Figure 2d); the TPA hydrogel showed a lower plateau (1,900–2,400 Pa) and crossover at 31–58% strain, consistent with dynamic bond dissipation in the reversible network.

Cyclic step-strain measurements confirmed injectability and self-recovery (Figure 2e). Under high strain the TFP hydrogel G′ dropped to ∼4 Pa (G″ > G′), indicating complete disruption into a flowable sol; upon strain removal, gel character was rapidly restored with G′ stabilizing near 500–518 Pa, reproducibly over four consecutive cycles. We note that this recovered modulus is intentionally measured after destructive strain and is therefore substantially lower than the modulus of the pristine, undisrupted network (Figure 2c, d); because β-ketoenamine crosslinks are irreversible covalent bonds, crosslinks broken during the high-strain step cannot re-form, and the partial yet robust recovery is itself direct evidence of the covalent nature of the network. All rheological measurements reported here were performed under identical buffer and temperature conditions. The hydrogel could be extruded through a G25 needle, highlighting its moldability and shape-retention capacity (Figure 2f).

### Load-Bearing Capacity and Shape Memory

To demonstrate the load-bearing capacity, we molded a wire-shaped object from the TFP hydrogel, which sustained a static tensile load of 900 g without fracture (Figure 3a), confirming the robustness of the β-ketoenamine network. The wires further exhibited reversible shape recovery on dehydration and rehydration (Figure 3b): on ambient drying, the wire underwent ∼21% length reduction (5.6 to 4.4 cm) while retaining the characteristic red β-ketoenamine chromophore, and water immersion quantitatively restored the original dimensions (5.9 cm, ∼105%); this cycle was fully reversible over four cycles without structural compromise. In contrast, conventional protein hydrogels based on physical or reversible imine crosslinks typically collapse irreversibly upon drying, underscoring the advantage of the COF-inspired strategy for dimensionally stable, processable materials.

### pH Stability

We evaluated pH stability by incubating gels in PBS spanning pH 1–13 at 37 °C, assessing integrity by tube inversion and visual inspection over 168 h (Figures 3c and S6). TFP hydrogels retained structural integrity and the characteristic red β-ketoenamine color at pH 4 –10 throughout, demonstrating exceptional stability across the physiologically relevant range; under extreme acidic (pH 1–3) and alkaline (pH 12–13) conditions, progressive color fading and partial disintegration appeared from 24 h onward, consistent with hydrolysis at extreme pH. TPA hydrogels, by contrast, dissolved rapidly, reflecting the hydrolytic lability of reversible imine crosslinks. Frequency-sweep rheology on gels incubated at pH 4, 5.5, 7.5, and 13 for 4 h (Figure 3d) showed G′ > G″ across the entire frequency range at all pH values, retaining elastic gel behaviour even at pH 13, where imine-based hydrogels dissolve completely. Storage moduli at 1 rad/s followed a pH-dependent trend, increasing from 453 Pa (pH 4) to 1,159 Pa (pH 5.5), 1,537 Pa (pH 7.5), and 9,041 Pa (pH 13). The low modulus at pH 4 reflects proximity to the isoelectric point of BSA (∼4.7), where reduced surface charge weakens electrostatic contributions, while the high modulus at pH 13 is attributed to partial alkaline unfolding that exposes additional lysine residues and increases crosslink density. The pristine-gel modulus in Figure 2c (∼33,500 Pa) is higher than these values because those measurements were performed on freshly formed gels in the alkaline carbonate gelation buffer, before equilibration in PBS at the pH values tested here; this is stated explicitly to avoid confusion between the two measurement conditions.

### Enzymatic Biodegradability

We assessed biodegradability by incubating TFP hydrogels in 1% Proteinase K solution at 37 °C, monitoring integrity by tube inversion over seven days (Figure 4a). The hydrogels degraded gradually over the seven days while protease-free controls remained intact, confirming enzyme-mediated rather than passive breakdown. Degradation over days rather than hours reflects the kinetic stability of the dense β-ketoenamine network against protease activity, an attractive feature for sustained-delivery applications.

**Figure 4.**
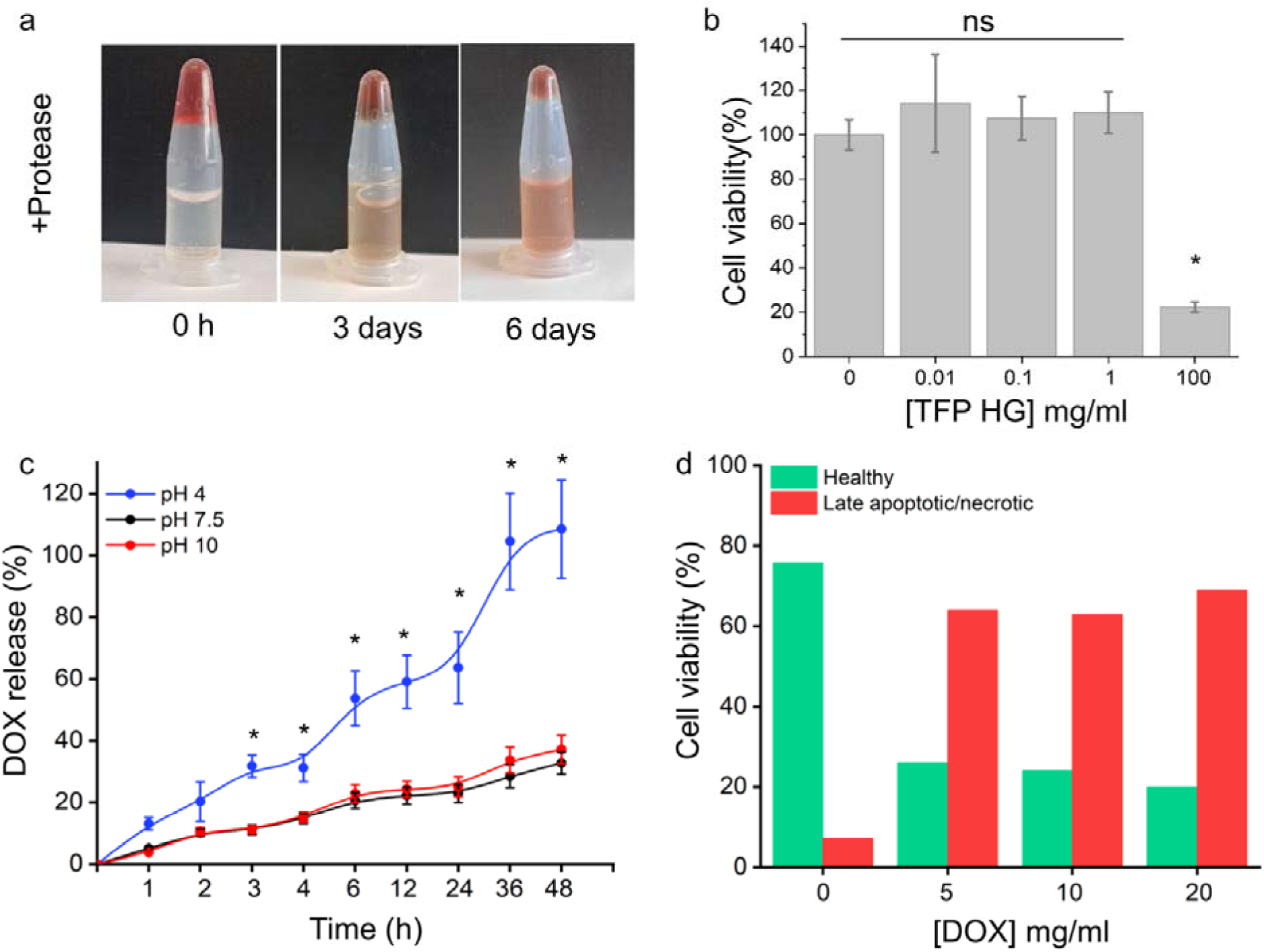
(a) Photographs of TFP hydrogels incubated with 1% Proteinase K at 0, 3, and 6 days, showing progressive enzyme-mediated degradation. (b) WST-1 viability of MCF-7 cells treated with the TFP hydrogel at 0, 0.01, 0.1, 1, and 100 mg/mL (mean ± SD, n = 3; ns = not significant; *p < 0.05 versus untreated control). (c) Cumulative pH-responsive DOX release from the TFP hydrogel at pH 4, 7.4, and 10 over 48 h (mean ± SD, n = 3; *p < 0.05). (d) Healthy and late-apoptotic/necrotic populations of MCF-7 cells determined by Annexin V/PI flow cytometry after 6 h treatment with DOX-loaded TFP hydrogel at DOX loading concentrations of 0, 5, 10, and 20 mg/mL.

### Drug Loading Capacity

Because the amount of drug introduced is never fully encapsulated, we quantified drug loading directly. A calibration curve constructed from free-DOX standards (0.5, 1, 1.5, 1.75, and 2 mg/mL) by intrinsic fluorescence gave the linear regression y = 2,924,095.5x + 4,737,304.4. Hydrogels prepared with nominal DOX loading concentrations of 0.5, 1.0, 1.5, and 1.75 mg/mL were measured by the same method, and the true encapsulated DOX concentration was back-calculated (Table S2). At low nominal loading (0.5 mg/mL), essentially all of the added DOX was retained within the hydrogel network (DLE ≈ 100%). As the nominal loading concentration increased, the encapsulated DOX concentration plateaued at approximately 1.09 mg/mL, and DLE fell to ∼62–72% at nominal loadings of 1.5–1.75 mg/mL. This plateau indicates a finite DOX encapsulation capacity under these conditions, consistent with DOX loading being governed by a limited number of available BSA drug-binding sites (Sudlow sites I and II) rather than by simple physical entrapment, which would be expected to scale linearly with the amount added.

### pH-Responsive Drug Release and Mechanism

We next investigated pH-responsive release of doxorubicin (DOX) from the TFP hydrogel by intrinsic fluorescence at pH 4 (acidic tumour microenvironment), 7.4 (physiological), and 10 (alkaline) over 48 h. DOX was loaded *in situ* before gelation, and fluorescence microscopy confirmed encapsulation (Figure S7). At pH 4, burst release reached 53.7 ± 8.9% at 6 h and 63.6 ± 11.5% at 24 h, with essentially quantitative release by 36–48 h. Release at pH 7.4 was significantly slower (22.9 ± 2.9% at 24 h; 32.8 ± 3.6% at 48 h), giving 2.8-fold selectivity for acidic over physiological pH at 24 h; release at pH 10 mirrored the pH 7.4 profile (25.1 ± 3.2% at 24 h; 37.2 ± 4.6% at 48 h), confirming acid-specific triggering rather than a general pH effect (Figure 4c).

To identify the release mechanism, we compared the release timescale against the independently measured timescales of network degradation and swelling rather than fitting a diffusion-based (Korsmeyer–Peppas) model, since our data argue against a simple diffusion/swelling-controlled process. First, degradation-controlled release can be excluded on kinetic grounds: the enzymatic degradation study (Figure 4a) shows that TFP hydrogels require Proteinase K and degrade gradually over seven days, while protease-free controls in buffer remain structurally intact for at least 168 h across pH 4-10 (Figure 3c). Because the DOX release experiments were conducted in protease-free PBS and were essentially complete within 36-48 h, the network is not undergoing meaningful mass loss or breakdown on the release timescale, ruling out bulk degradation as the driver. Second, swelling alone cannot account for the magnitude of the pH selectivity: the equilibrium swelling ratios measured above show only a modest difference between pH 5.5 (Q = 1.19) and pH 7.4 (Q = 1.17), a ∼2% difference in water uptake far too small to explain the 2.8-fold difference in cumulative release observed between pH 4 and pH 7.4 at 24 h. Third, release is instead attributed principally to a pH-dependent decrease in DOX–BSA binding affinity: the N-to-F conformational transition of BSA near its isoelectric point (∼4.7) is known to weaken hydrophobic drug–protein interactions at Sudlow sites I and II specifically under acidic conditions, independent of bulk network swelling. This is consistent with the sharp, acid-selective release profile that closely tracks pH rather than modest swelling differences, and with the similarity between the pH 7.4 and pH 10 release profiles, both far from the isoelectric point and both showing similarly low release. In contrast, only pH 4, close to the isoelectric point, triggers burst release. We therefore conclude that pH-responsive DOX release is governed primarily by pH-dependent conformational weakening of DOX–BSA binding, with a secondary, minor contribution from acid-enhanced network swelling, while bulk hydrogel degradation is excluded as a contributing mechanism on the 24-48 h release timescale.

A different release pattern emerged for ciprofloxacin (Cipro) at pH 7.4 (Figure S8). Rather than the sharp, pH-triggered burst seen for DOX, cipro was released gradually and continuously throughout the 48 h window around 20% released from the network by 6 h, climbing to about 38% by 10–12 h. Release then slowed considerably, hovering near 51–53% from 18 to 30 h, a plateau that might have suggested near-complete release, were it not for a subsequent rise to 73 ± 10% by 48 h. This two-phase behaviour, an early diffusion-like rise followed by a slower secondary release, points to a mechanism quite unlike the one governing DOX. Cipro is small, hydrophilic, and zwitterionic, and its interactions with the BSA-containing network are unlikely to involve the same conformationally gated, site-specific binding (Sudlow sites I/II) that controls DOX release. Instead, the data are more consistent with diffusion through the swollen network dominating early release, with the later-stage increase potentially reflecting slow network relaxation or a secondary, weaker binding population.

### In Vitro Anticancer Efficacy and Cytocompatibility

We assessed the cytocompatibility of the drug-free TFP hydrogel against MCF-7 cells by WST-1 assay (Figure 4b). At 0.01, 0.1, and 1 mg/mL, viability was 114.2 ± 22.0%, 107.6 ± 9.8%, and 110.0 ± 9.3%, respectively, statistically indistinguishable from the untreated control (100 ± 6.9%), confirming the absence of cytotoxicity across this range; the mild enhancement is consistent with the growth-supporting properties of BSA.^33^ At 100 mg/mL, viability decreased to 22.3 ± 2.3%. We emphasize that this concentration is far above the intended working range: the material functions as a localized, injectable depot rather than being dispersed at high concentration in the cellular environment, and the therapeutic effect derives from released DOX rather than from matrix toxicity. The excellent biocompatibility at ≤1 mg/mL reflects the chemical stability of the β-ketoenamine crosslinks, which precludes leaching of toxic crosslinker residues, a key advantage over glutaraldehyde-based systems, where residual-crosslinker release is a persistent concern.^19,20^ Cytocompatibility against the normal cell line HEK 293 gave 80 % viability at 20 mg/mL, establishing the biosafety of the matrix (Figure S9).

To evaluate DOX efficacy against MCF-7 cells, we performed Annexin V/PI flow cytometry after 6 h of treatment with DOX-loaded TFP hydrogels, presenting the quantified healthy and late-apoptotic/necrotic populations in Figure 4d (representative dot plots in Figure S10). Free DOX (1.5 µg/mL) reduced viability to 49.8% with 35.6% apoptotic or necrotic cells. In contrast, DOX-loaded TFP hydrogels at 5, 10, and 20 mg/mL induced 64%, 63%, and 69% apoptosis and necrosis, with healthy populations of only 26%, 24%, and 20%, representing ∼1.8-fold higher cell death than free DOX at the lowest loading. We attribute the enhanced killing to pH-triggered burst release combined with albumin-receptor-mediated (gp60/SPARC) internalization of BSA–DOX complexes by MCF-7 cells.^34^

### Antibacterial Activity of Ciprofloxacin-Loaded Hydrogel (HG+Cipro)

Bacterial infections pose a major threat in biomedical applications, particularly in wound healing, tissue engineering, and implantable biomaterials, where infection can impede regeneration and lead to treatment failure^35^. To address this, we developed a ciprofloxacin-loaded hydrogel (HG+Cipro) and assessed its antibacterial efficacy against both Gram-positive (Micrococcus luteus) and Gram-negative (Escherichia coli) strains using multiple assays. We checked the antibacterial effect of the ciprofloxacin-loaded hydrogel (HG+Cipro) against E. coli and *M. luteus* using three approaches, i.e., bacterial zone inhibition test, in vitro surface antibacterial test, and antibacterial growth kinetics test^36^.

The drug-loaded gel produced comparable zones of inhibition against both species, with diameters of 43.87 ± 1.65 mm against E. coli and 43.71 ± 2.92 mm against *M. luteus* (Figure 5b), indicating no strong species selectivity in this assay. The slow diffusion of ciprofloxacin from the gel inhibits bacterial cell multiplication, resulting in zone formation. Gram-negative bacteria such as E. coli possess an outer lipopolysaccharide membrane that Gram-positive bacteria such as *M. luteus* lack; this additional barrier can restrict antibiotic access in Gram-negative species, so the similar zone sizes observed here indicate that ciprofloxacin diffusing from the hydrogel was able to penetrate both cell envelopes effectively. Surface-treated bacteria were spread on the agar plate and incubated for 12–16 hours (Figure 5a). It was observed that the control and the hydrogel sample exhibited normal bacterial growth, while the cipro-loaded hydrogel showed no growth on the plate. A kinetic study (Figure 5c) of bacterial growth was also performed for different hydrogel combinations for 12 hours at an interval of 2 hours^37^. It was noticed that the HG+Cipro combination had an extended lag phase compared to HG and control due to the antibacterial action of the drug killing most of the bacteria, while the control group had intact cell wall. A large amount of debris and cell wall damage was observed in the treated (Hg-Cipro) group. This strong effect comes from ciprofloxacin’s action of blocking DNA gyrase and topoisomerase IV, enzymes that bacteria need to multiply. Overall, our results show that HG+Cipro can strongly inhibit both Gram-positive and Gram-negative bacteria, making it a promising material for preventing infections in wound healing, implants, and other biomedical uses.

**Figure 5.**
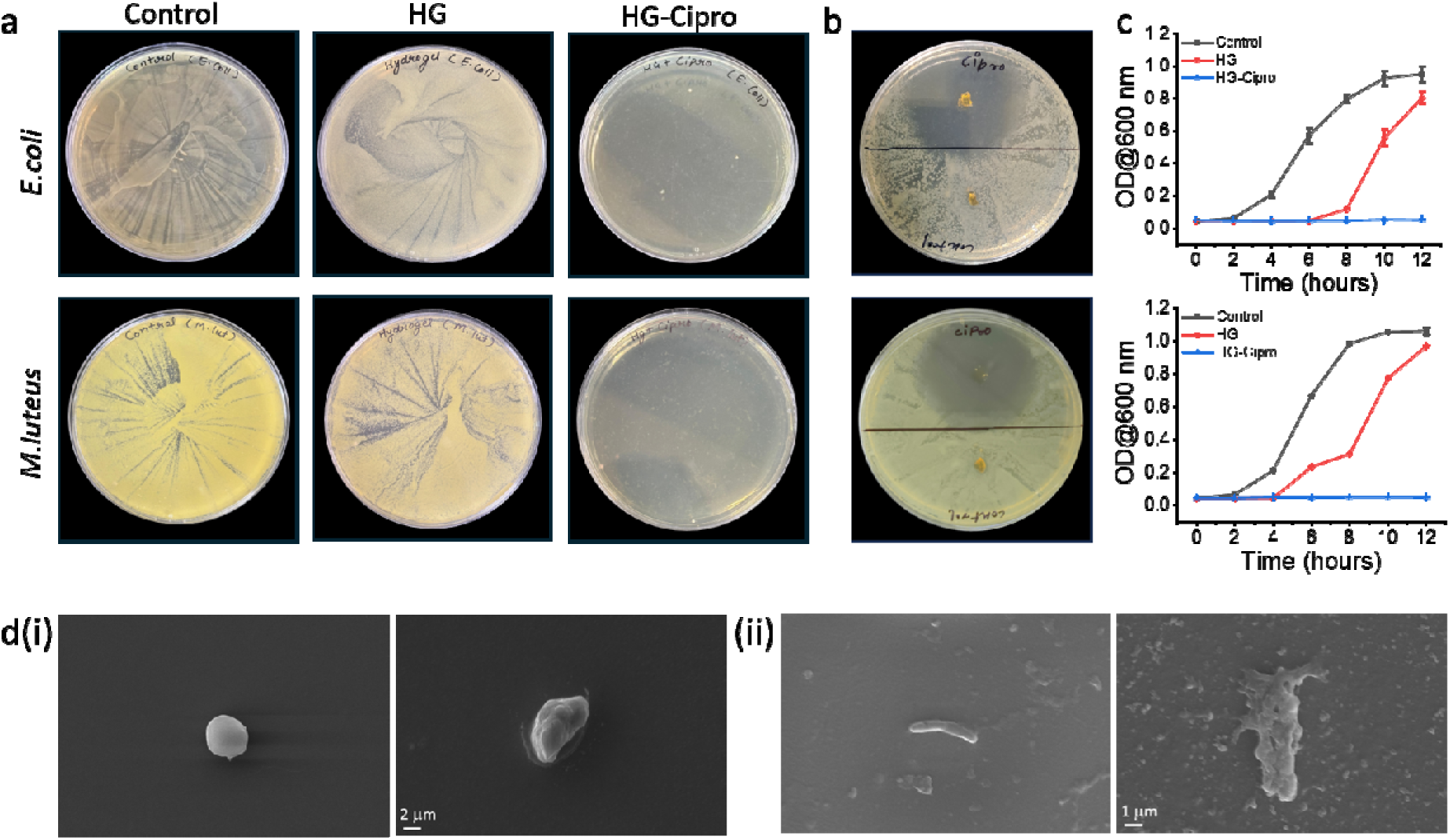
Antibacterial effect of hydrogel (HG) and HG-Ciprofloxacin (HG+Cipro) on *E. coli* and *M. luteus*. (a) Streak plate assays showing bacterial growth on Control, HG, and HG+Cipro treated plates for *E. coli* (top) and *M. luteus* (bottom). (b) Zone-of-inhibition assays comparing Cipro and hydrogel-treated samples against the two bacterial strains. (c) Bacterial growth kinetics (OD@600 nm) over 12 h for Control, HG, and HG+Cipro groups against *E. coli* (top) and *M. luteus* (bottom), showing significant growth inhibition in the HG+Cipro group. (d) SEM images of (i) untreated bacteria displaying intact, smooth morphology (scale bar: 2 µm) and (ii) HG+Cipro-treated bacteria showing pronounced surface damage and structural deformation (scale bar: 1 µm).

## CONCLUSIONS

We have demonstrated that the irreversible β-ketoenamine bond-forming chemistry that underpins keto-enamine covalent organic frameworks can be transferred to a native globular protein, crosslinking the surface lysine residues of BSA with the trialdehyde TFP into a chemically defined hydrogel under mild aqueous conditions and without denaturation. FTIR and NMR establish the irreversible β-ketoenamine linkage, and tryptophan fluorescence confirms retention of the native fold. The resulting hydrogel uniquely combines high stiffness (G′ ≈ 33,500 Pa, ∼40-fold above the reversible-imine control), injectability and self-recovery, reversible shape memory, a 900 g load-bearing capacity, broad pH stability (pH 4–10, 168 h), and enzymatic degradability within a single, structurally defined network. The hydrogel additionally reaches an equilibrium swelling ratio of Q ≈ 1.17 at physiological pH, adopts a dense, nodular network morphology by SEM relative to the more open, lamellar reversible-imine control, and encapsulates doxorubicin with a loading efficiency of up to ∼100% at low input, plateauing near 1.09 mg/mL of encapsulated drug at higher loadings. The hydrogel delivers doxorubicin in a pH-triggered, acid-selective manner governed principally by pH-dependent weakening of DOX–BSA binding affinity rather than by bulk swelling or degradation, producing enhanced killing of MCF-7 cells. A ciprofloxacin-loaded variant of the hydrogel further exhibits strong antibacterial activity against both Gram-negative (E. coli) and Gram-positive (M. luteus) bacteria, pointing to a role for this platform in infection-resistant wound-healing and implant coatings alongside its drug-delivery function. More broadly, transplanting irreversible β-ketoenamine chemistry from crystalline frameworks to soft biological matter offers a general route to mechanically robust, stimuli-responsive protein biomaterials, and we anticipate its extension to other proteins, enzymes, and growth factors for tissue engineering and on-demand drug delivery.

## ASSOCIATED CONTENT

### Supporting Information

The Supporting Information is available free of charge at https://pubs.acs.org/doi/XXXXX. Materials and full experimental procedures; FTIR spectra of crosslinking chemistry (Figures S1, S2); solution-state NMR spectra (Figure S3); tryptophan fluorescence spectra (Figure S4); SEM micrographs (Figure S5); pH-stability photographs (Figure S6); DOX fluorescence micrograph (Figure S7), ciprofloxacin release profile (Figure S8); cytotoxicity in HEK 293 cells (Figure S9), representative Annexin V/PI flow-cytometry dot plots (Figure S10); swelling data (Table S1); drug loading capacity/efficiency data (Table S2).

## AUTHOR INFORMATION

### Corresponding Authors

Suchetan Pal - Department of Chemistry and Department of Bioscience and Biomedical Engineering, Indian Institute of Technology Bhilai, Durg 491002, India; ORCID: 0000-0002-6323-7502;

Tatini Rakshit - Department of Chemistry, Shiv Nadar Institution of Eminence, Greater Noida 201314, India;

### Notes

The authors declare no competing financial interest.

## ACKNOWLEDGMENTS

S.P. thanks the Science and Engineering Research Board (CRG/2023/003029) and the Anusandhan National Research Foundation (ANRF/ARG/2025/005072/LS). The authors thank UGC, MHRD, the Central Instrumentation Facility, and the Department of Chemistry, Indian Institute of Technology Bhilai, and acknowledge the Shiv Nadar Institution of Eminence for rheometer facilities used in this study.

