## Supplemental file for "A Covalent Organic Framework-Inspired β-Ketoenamine Crosslinking Strategy for Robust, Injectable Bovine Serum Albumin Hydrogels with pH-Triggered Drug Release"

**A Covalent Organic Framework-Inspired β-Ketoenamine-Crosslinked Bovine Serum Albumin Hydrogel for pH-Triggered Anticancer and Antibacterial Drug Delivery**

Sonal Khaitan,^a^ Tanya Agrawal,^b^ Parth Gulati,^c^ Akansha,^a^ Natasha,^b^ Tatini Rakshit, ^*b^ and Suchetan Pal^*a,c^

*^a^ Department of Chemistry, Indian Institute of Technology Bhilai, Durg 491002, Chhattisgarh, India*

*^b^ Department of Chemistry, Shiv Nadar Institution of Eminence, Greater Noida 201314, Uttar Pradesh, India*

*^c^ Department of Bioscience and Biomedical Engineering, Indian Institute of Technology Bhilai, Durg 491002, Chhattisgarh, India*

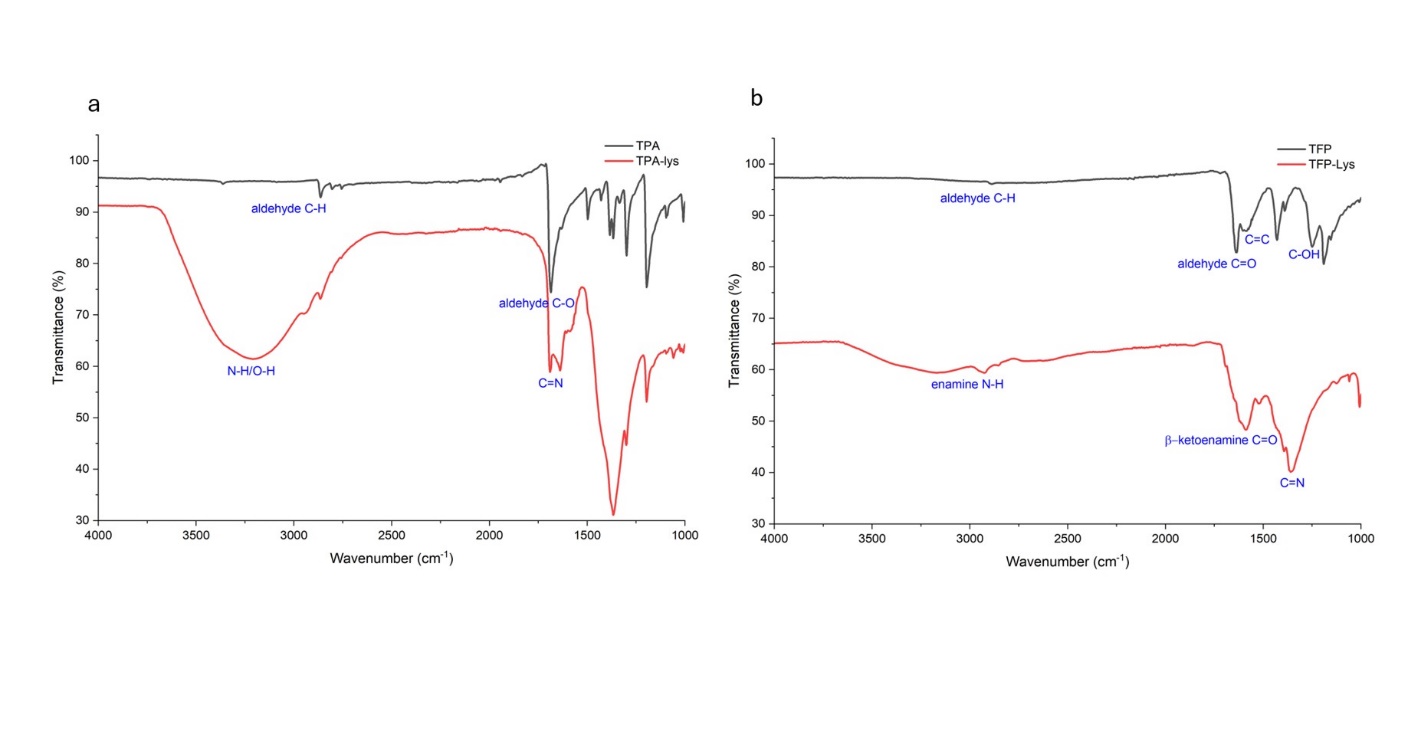
Fig. S1. FTIR spectrum of (a) TPA-Lys (red) and TPA (black) (b) TFP-Lys (red) and TFP (black).

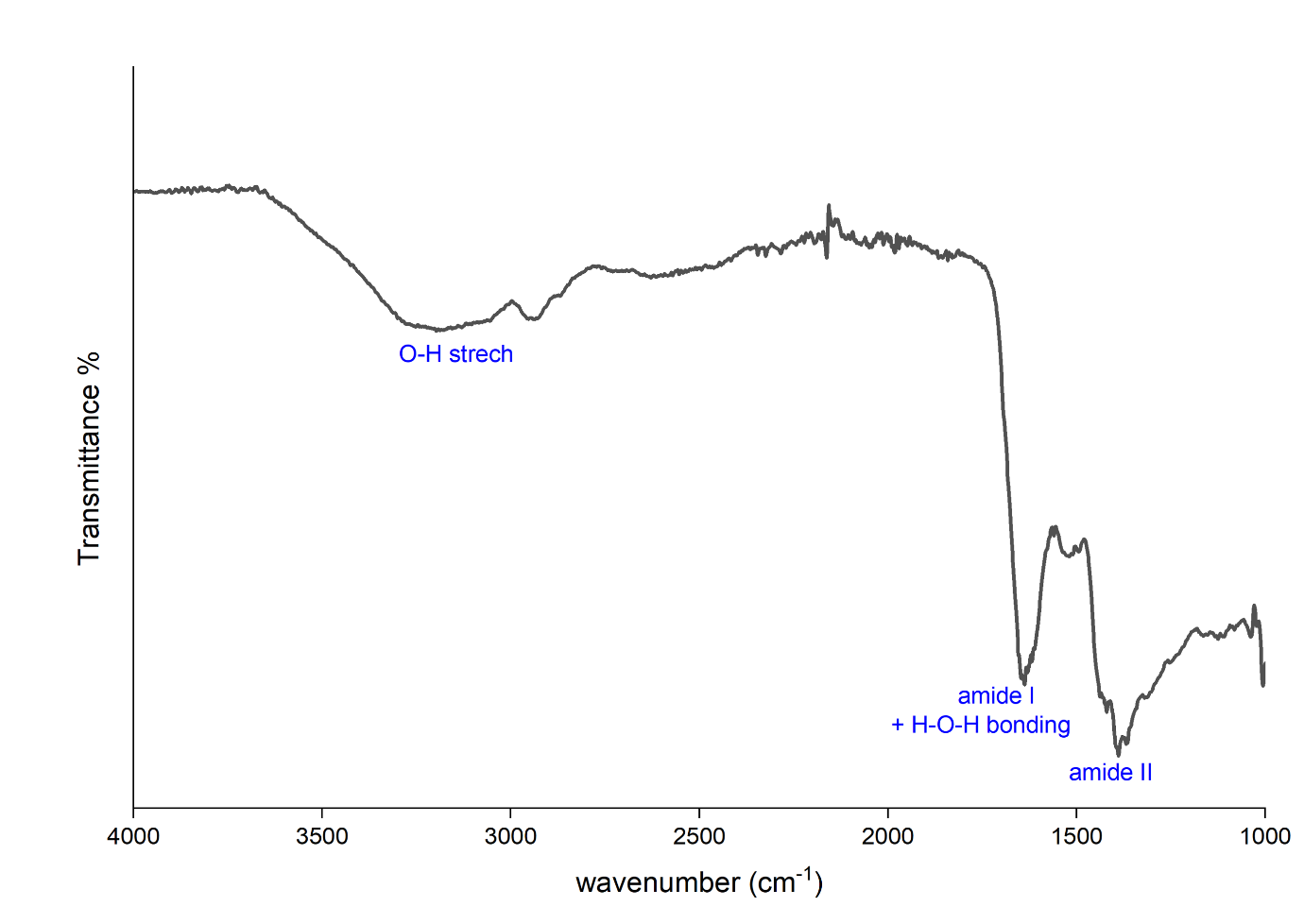

Fig. S2. FTIR spectrum of TFP crosslinked BSA hydrogel.

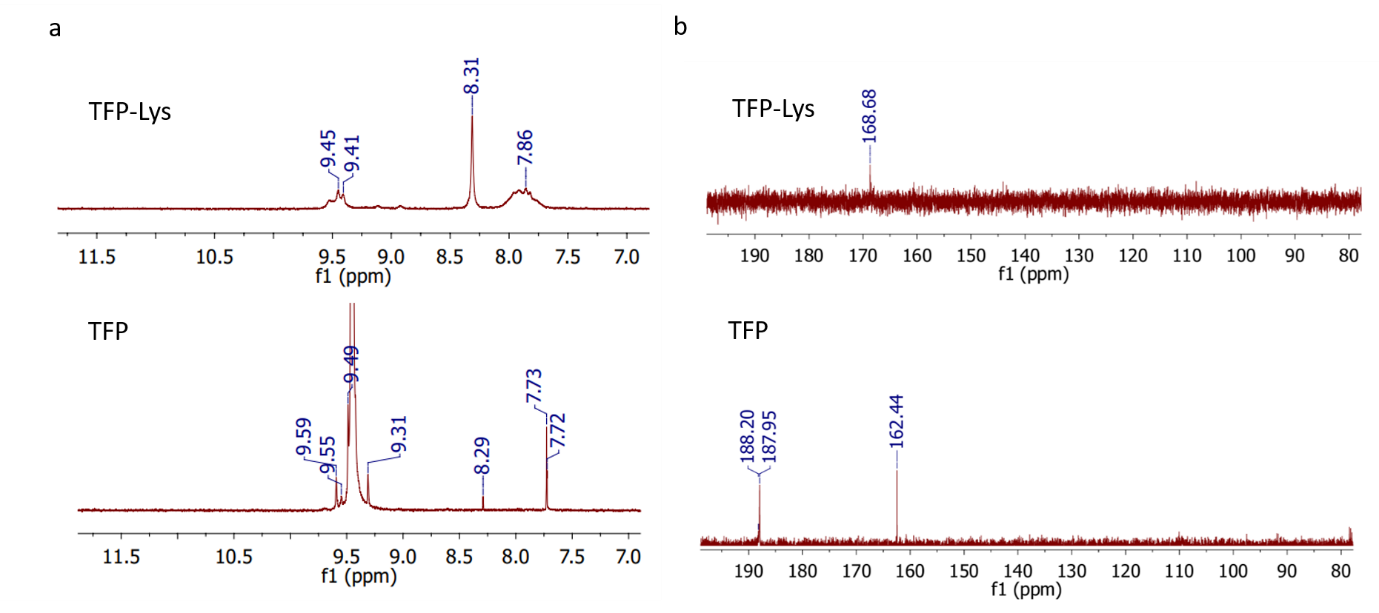

Fig. S3. (a) ^1^H NMR peaks of TFP-Lys and TFP. (b) ^13^C NMR peaks of TFP-Lys and TFP.

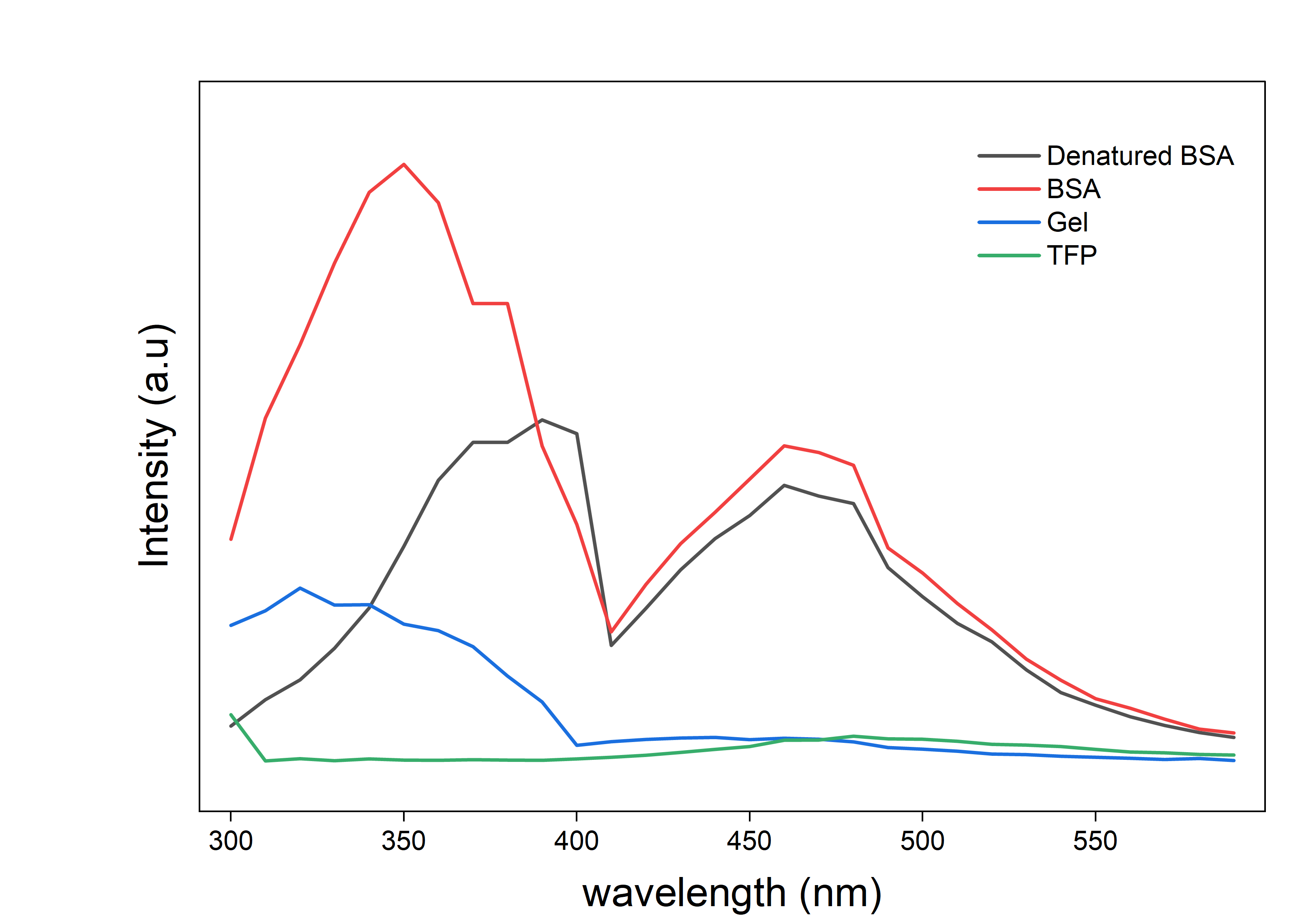

Fig. S4. Fluorescence emission spectra of native BSA (red), denatured BSA (black), BSA-TFP hydrogels (blue), and TFP only (green).

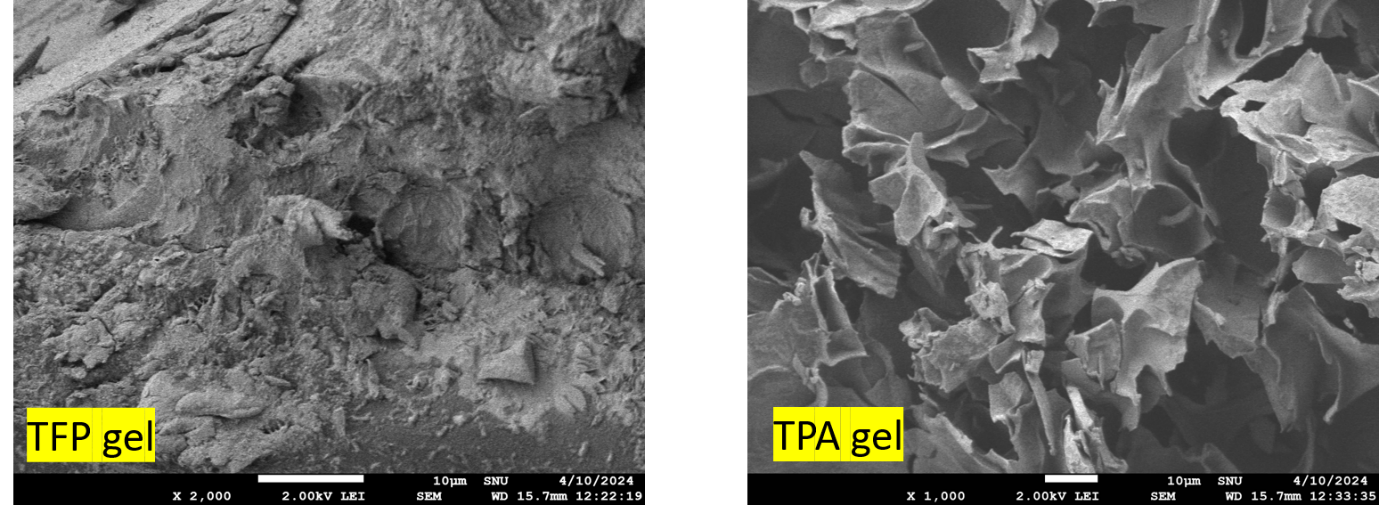

Fig. S5. SEM micrographs of TFP and TPA hydrogels.

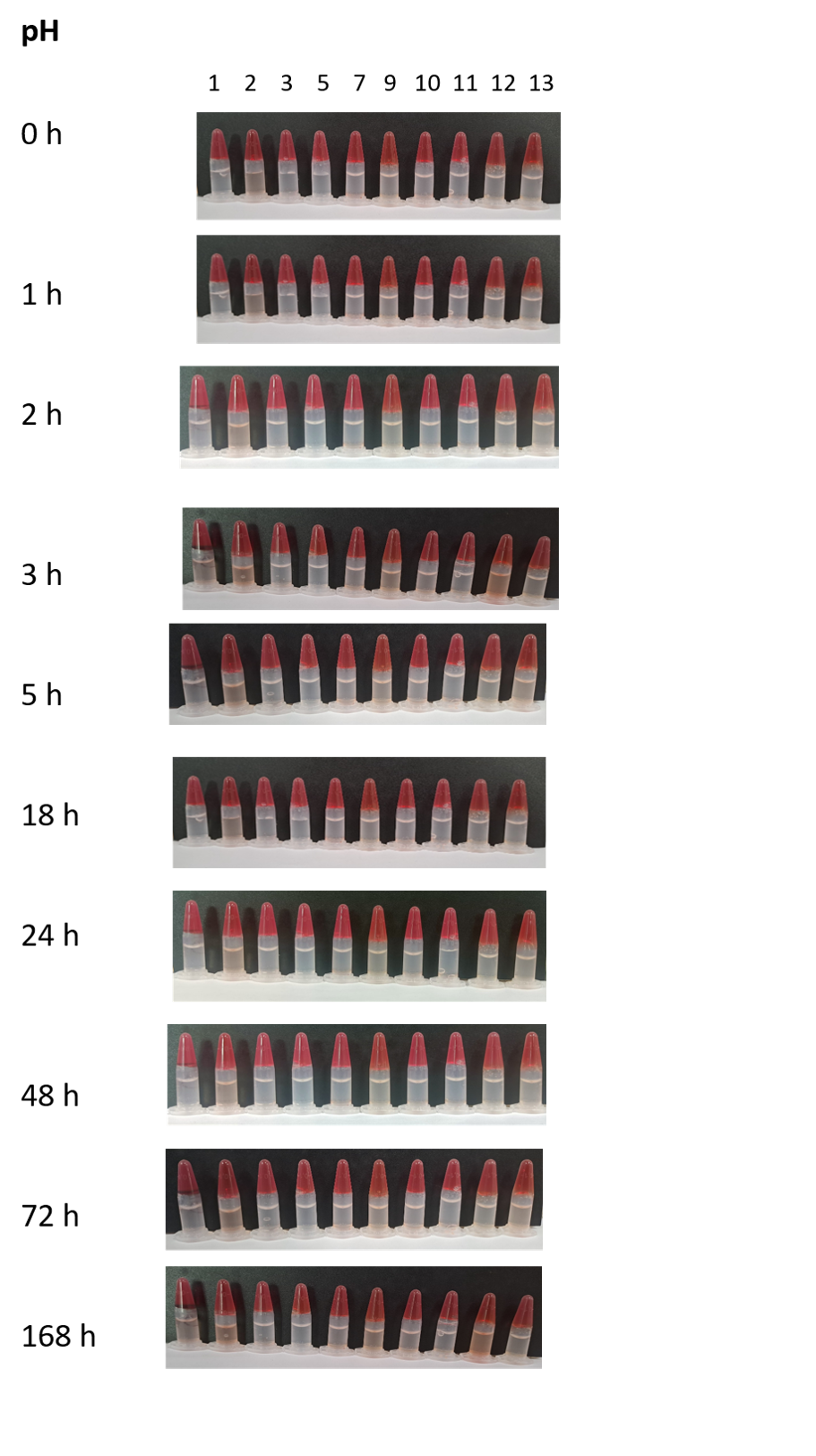

Fig. S6. Photographs of TFP HGs in PBS at pH 1, 2, 3, 5, 7, 9, 10, 11, 12, 13 at different points of incubation.

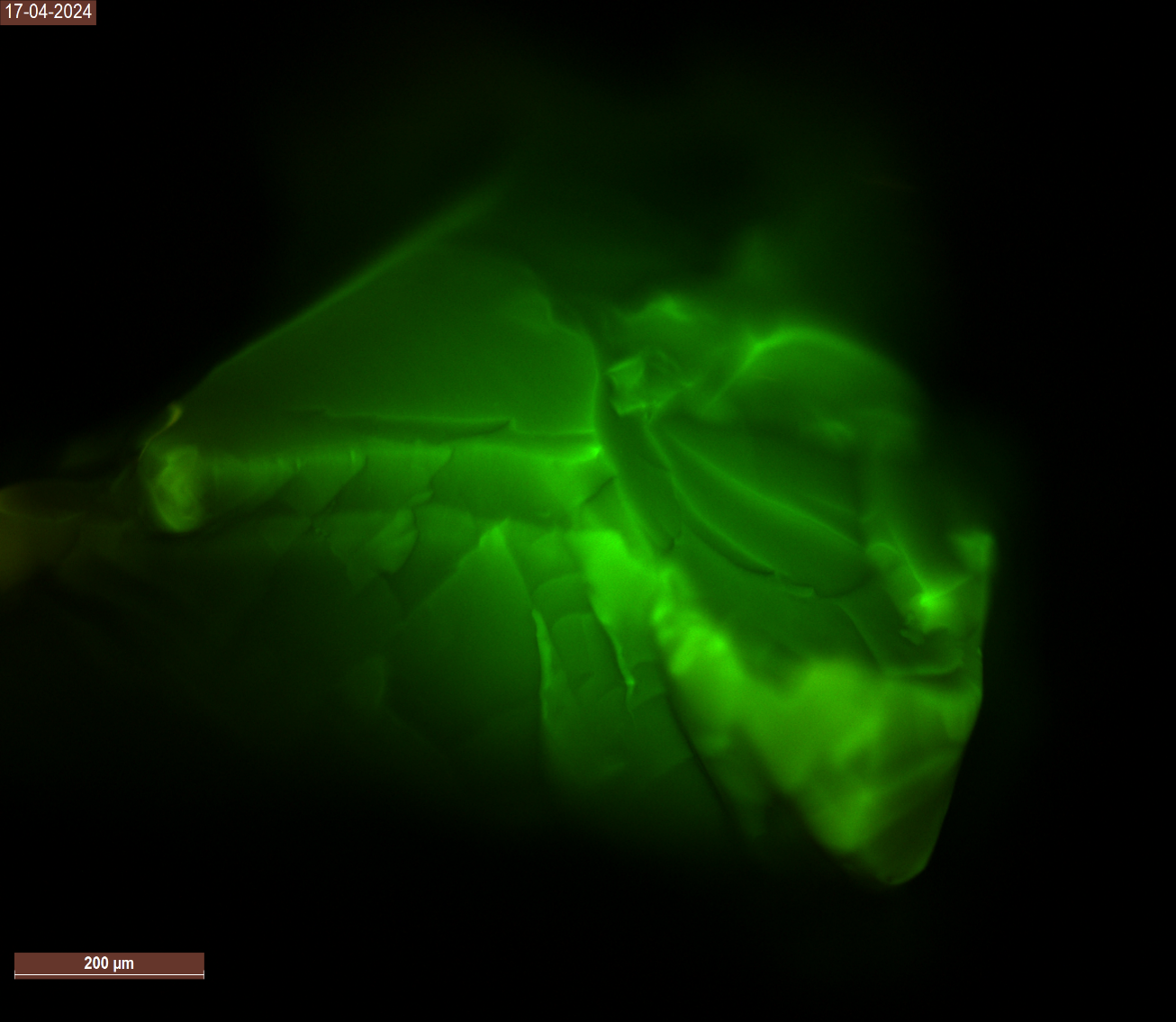

Fig. S7. Fluorescence micrograph of DOX-loaded TFP HG (scale bar 200 μm).

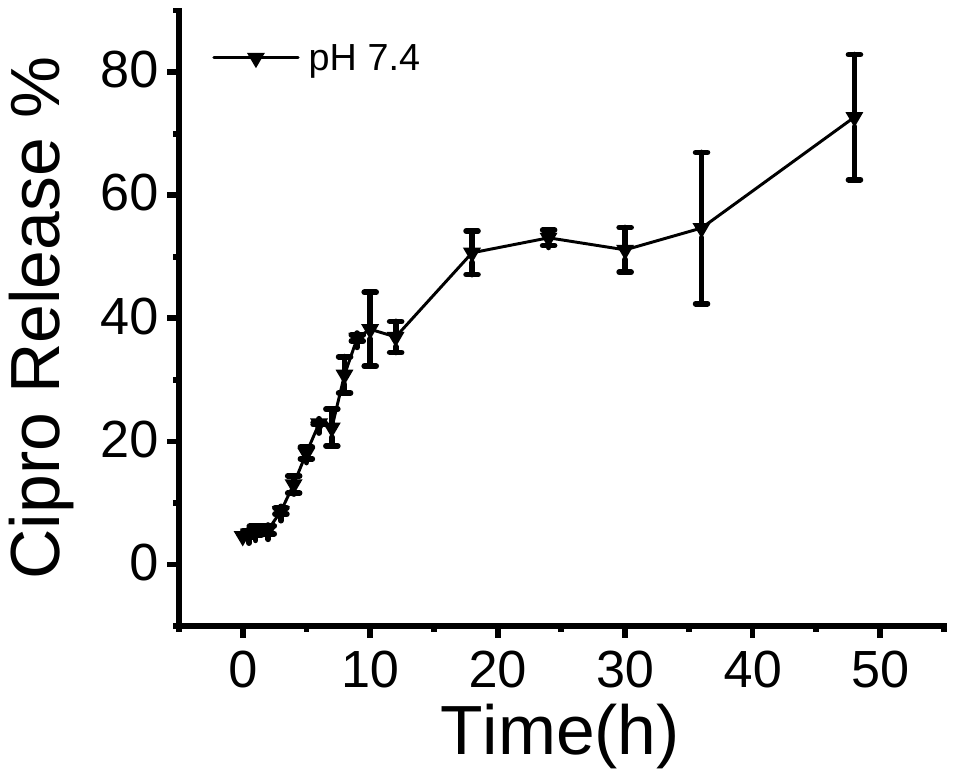

Fig. S8. Cumulative release (%) of ciprofloxacin from the hydrogel.

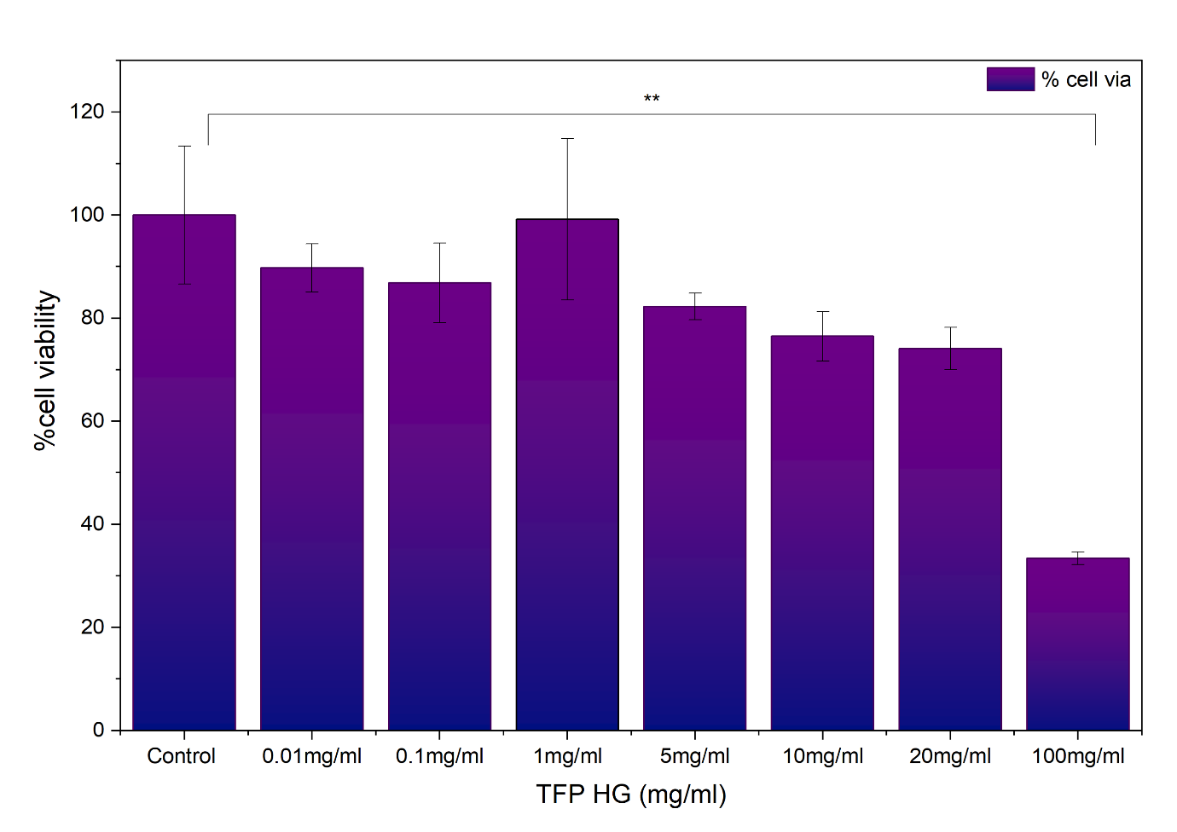

Fig. S9. WST-1 cytotoxicity assay of TFP HG against HEK293 cells.

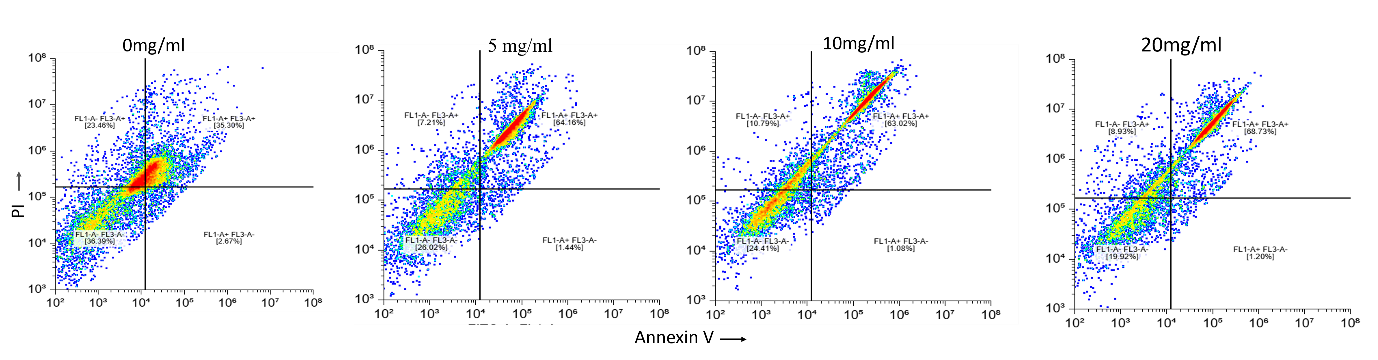

Fig. S10. Flow cytometry data (dot plot) showing DOX-induced cytotoxicity in MCF-7 Cells.

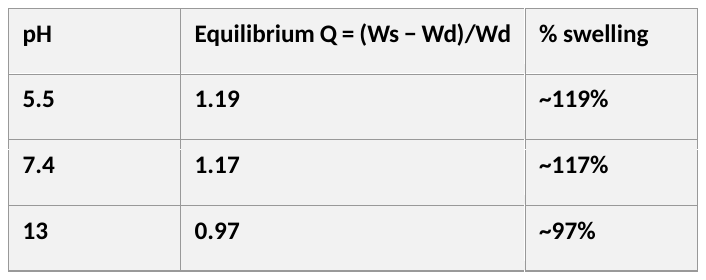

Table S1. Equilibrium (24 h) swelling ratio of the TFP hydrogel as a function of pH.

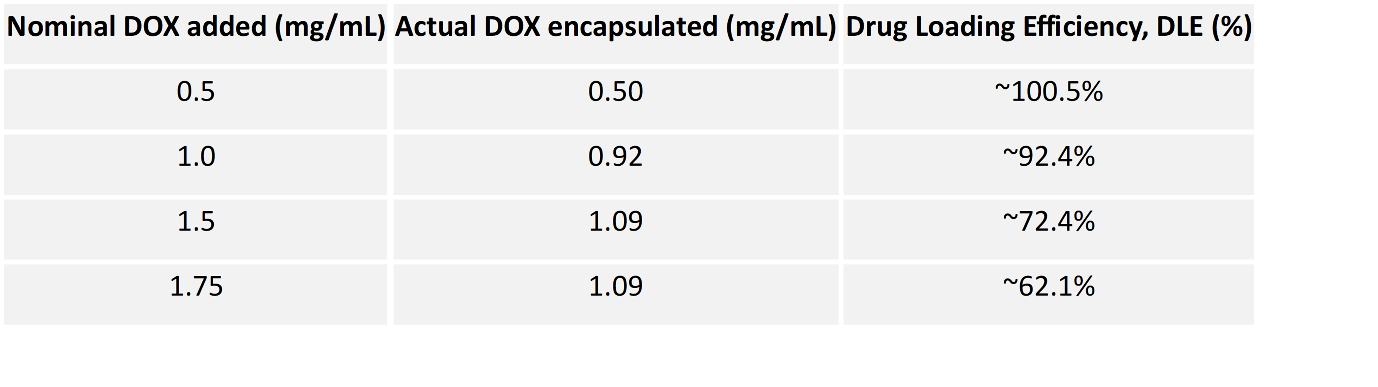

Table S2. Drug loading efficiency (DLE) of the TFP hydrogel as a function of nominal DOX input.
